# Tetranucleotide frequencies differentiate genomic boundaries and metabolic strategies across environmental microbiomes

**DOI:** 10.1101/2025.01.16.633425

**Authors:** Matthew Kellom, Maureen Berg, I-Min A. Chen, Ken Chu, Alicia Clum, Marcel Huntemann, Natalia N. Ivanova, Nikos C. Kyrpides, Supratim Mukherjee, T. B. K. Reddy, Simon Roux, Rekha Seshadri, Gitta Szabo, Neha J. Varghese, Tanja Woyke, Emiley A. Eloe-Fadrosh

## Abstract

Microbiomes are constrained by physicochemical conditions, nutrient regimes, and community interactions across diverse environments, yet genomic signatures of this adaptation remain unclear. Metagenome sequencing is a powerful technique to analyze genomic content in the context of natural environments, establishing concepts of microbial ecological trends. Here, we developed a data discovery tool - a tetranucleotide-informed metagenome stability diagram - that is publicly available in the Integrated Microbial Genomes and Microbiomes (IMG/M) platform for metagenome-ecosystem analyses. We analyzed the tetranucleotide frequencies from quality-filtered and unassembled sequence data of over 12,000 metagenomes to assess ecosystem-specific microbial community composition and function. We found that tetranucleotide frequencies can differentiate communities across various natural environments, and that specific functional and metabolic trends can be observed in this structuring. Our tool places metagenomes sampled from diverse environments into clusters and along gradients of tetranucleotide frequency similarity, suggesting microbiome community compositions specific to gradient conditions. Within the resulting metagenome clusters, we identify protein-coding gene identifiers that are most differentiated between ecosystem classifications. We plan for annual updates to the metagenome stability diagram in IMG/M with new data, allowing for refinement of the ecosystem classifications delineated here. This framework has the potential to inform future studies on microbiome engineering, bioremediation, and the prediction of microbial community responses to environmental change.

**Importance:** Microbes adapt to diverse environments influenced by factors like temperature, acidity, and nutrient availability. We developed a new tool to analyze and visualize the genetic makeup of over 12,000 microbial communities, revealing patterns linked to specific functions and metabolic processes. This tool groups similar microbial communities and identifies characteristic genes within environments. By continually updating this tool, we aim to advance our understanding of microbial ecology, enabling applications like microbial engineering, bioremediation, and predicting responses to environmental change.

## Introduction

Throughout the course of Earth’s history, microbial populations have adapted to survive and grow in a wide range of environmental conditions. Every environment hosts a combination of physicochemical conditions, nutrient regimes, and community interactions that shape the fitness of specific taxa and their evolved metabolisms (Hamilton and Havig 2022; Hamilton et al. 2019; Gutiérrez-Preciado et al. 2018; Vigneron et al. 2022; Bernhardt et al. 2020). The diversity of these adaptations is captured in the genomes of individual microbes that coalesce into environment-specific microbiomes, suited to their conditions (J. M. Dick et al. 2023). This means that environmental metagenomes collectively represent the genomic traits of microbial communities under a myriad of selective pressure gradients, and their genomes should reflect differences in those selective pressures (Yan et al. 2020; Glassman et al. 2018). In disturbance ecology, maintaining relatively constant alpha-diversity when microbial communities are exposed to selective pressures is referred to as microbiome stability (Pimm 1984; Sorensen and Shade 2020). It has been shown that similar environments harbor similar stable microbiomes (Raymond and Alsop 2015; Sriaporn et al. 2023; G. J. Dick et al. 2009), and stable microbiomes have been observed along environmental gradients (Cox, Shock, and Havig 2011; Alsop, Boyd, and Raymond 2014; Chase, Weihe, and Martiny 2021; Swingley et al. 2012; Glassman et al. 2018). However, the extent to which microbial populations and metabolic functions can be distributed across environmental conditions is often unclear, inhibiting our ability to predict population composition and metabolic activity from environmental context (Gibbons and Gilbert 2015; Glassman et al. 2018; Chase, Weihe, and Martiny 2021; Flinkstrom et al. 2024).

Shotgun metagenomics offers a lens through which to capture the taxonomic and putative functional diversity of microbial communities, enabling analyses of microbiome ecological trends across different environment types at a global scale (Schmidt, DeLong, and Pace 1991; Grossart et al. 2020; G. J. Dick et al. 2009). As metagenome databases continue to grow, meta-analyses can leverage data across thousands of disparate datasets (Nayfach et al. 2021; Roux et al. 2019; Chen et al. 2023; Richardson et al. 2023; Chikhi et al. 2024). Despite the massive increase in sequence data across diverse environments, one key barrier for deriving cross-environment ecological insights has been the sparseness of contextual information or metadata(Kyrpides, Eloe-Fadrosh, and Ivanova 2016; Huttenhower, Finn, and McHardy 2023; Eloe-Fadrosh et al. 2024). One source of metadata that is often required for data submission, and thus regularly associated with metagenome datasets, is environment ecosystem classification (Mukherjee et al. 2023). While broad ecosystem classifications may orient researchers toward major taxonomy and metabolic processes expected within a sample (Duan et al. 2022), the interplay between microbial communities and their environments is multifaceted, leading to a level of complexity that can make direct comparisons and predictions challenging (Gonze et al. 2017; VanInsberghe et al. 2020).

Previous studies have utilized diversity metrics or functional profiling to differentiate microbiome composition across ecosystems (Tringe et al. 2005; Dinsdale et al. 2008; Thompson et al. 2017; Flinkstrom et al. 2024). Here, we hypothesized that tetranucleotide frequencies from unassembled metagenome data could be used as a proxy to evaluate ecosystem-specific microbiome composition and applied this analysis to over 12,000 environmental metagenomes. Tetranucleotides, composed of four DNA bases, are universally shared components of metagenomes with usage patterns that have phylogenetic-resolving specificity not seen in di- or tri-nucleotides (Noble, Citek, and Ogunseitan 1998; Pride et al. 2003; Teeling et al. 2004). Their small size allows for a relatively low possibility of 256 combinations to compare between metagenomes, which can be further reduced to 136 when accounting for duplicate frequencies of reverse complements (Zhou, Olman, and Xu 2008). Converting tetranucleotide counts into frequencies (percentages relative to all tetranucleotide counts combined) normalizes usage patterns to be independent of sequencing data size, with some potential biases still present due to variations in sequencing depth and coverage of lower abundance taxa (Lynch and Neufeld 2015). We frame tetranucleotide frequencies as components of metagenomes, analogous to elements as components of minerals. In the geosciences, mineral stability diagrams depict two conditions for a given set of minerals: 1) axes that show physicochemical conditions necessary for minerals to coexist at equilibrium, and 2) which minerals are stable and which are unstable along those axes (Faure 1998). We group metagenomes with respect to sampling environment using the Genomes OnLine (GOLD) Ecosystem classifications (Mukherjee et al. 2023) and show that tetranucleotide frequencies can be used as a signature to map metagenome compositions in a tetranucleotide-informed metagenome stability diagram. We extracted ranked lists of gene annotations that displayed differential abundances across all ecosystem classifications, along with gene annotations that differentiated specific ecosystems. To facilitate future analyses, we have implemented the tetranucleotide-informed metagenome stability diagram as a service available on the Department of Energy (DOE) Joint Genome Institute’s (JGI) Integrated Microbial Genomes & Microbiomes (IMG/M) platform (https://img.jgi.doe.gov/cgi-bin/mer/main.cgi?section=AdvAnalytics&page=environmental) (Chen et al. 2023).

## Results

### Development of a tetranucleotide-informed metagenome stability diagram

Microbiomes from diverse ecosystems can be differentiated based on diversity metrics and unique functional profiles (Tringe et al. 2005; Dinsdale et al. 2008; Thompson et al. 2017; Flinkstrom et al. 2024). We leveraged tetranucleotide frequencies combined with linear discriminant analysis (LDA) dimensionality reduction and k-nearest neighbors (KNN) classification to generate a tetranucleotide-informed metagenome stability diagram for 12,063 metagenomes (**Fig. 1**). These metagenomes span diverse natural environments with the majority sampled from soil (5,236), marine (2,761), and freshwater (2,481) ecosystems and have been manually curated with the five-level GOLD ecosystem classification scheme (Mukherjee et al. 2023) (**Supplemental File S1**). We assumed that all naturally occurring metagenomes will contain all possible tetranucleotide combinations as universal basic shared components (Pride et al. 2003). Both tetranucleotide frequencies and ratios of tetranucleotide frequencies were used for metagenome analysis (**Supplemental File S2**), as described in the Methods.

**Figure 1.**
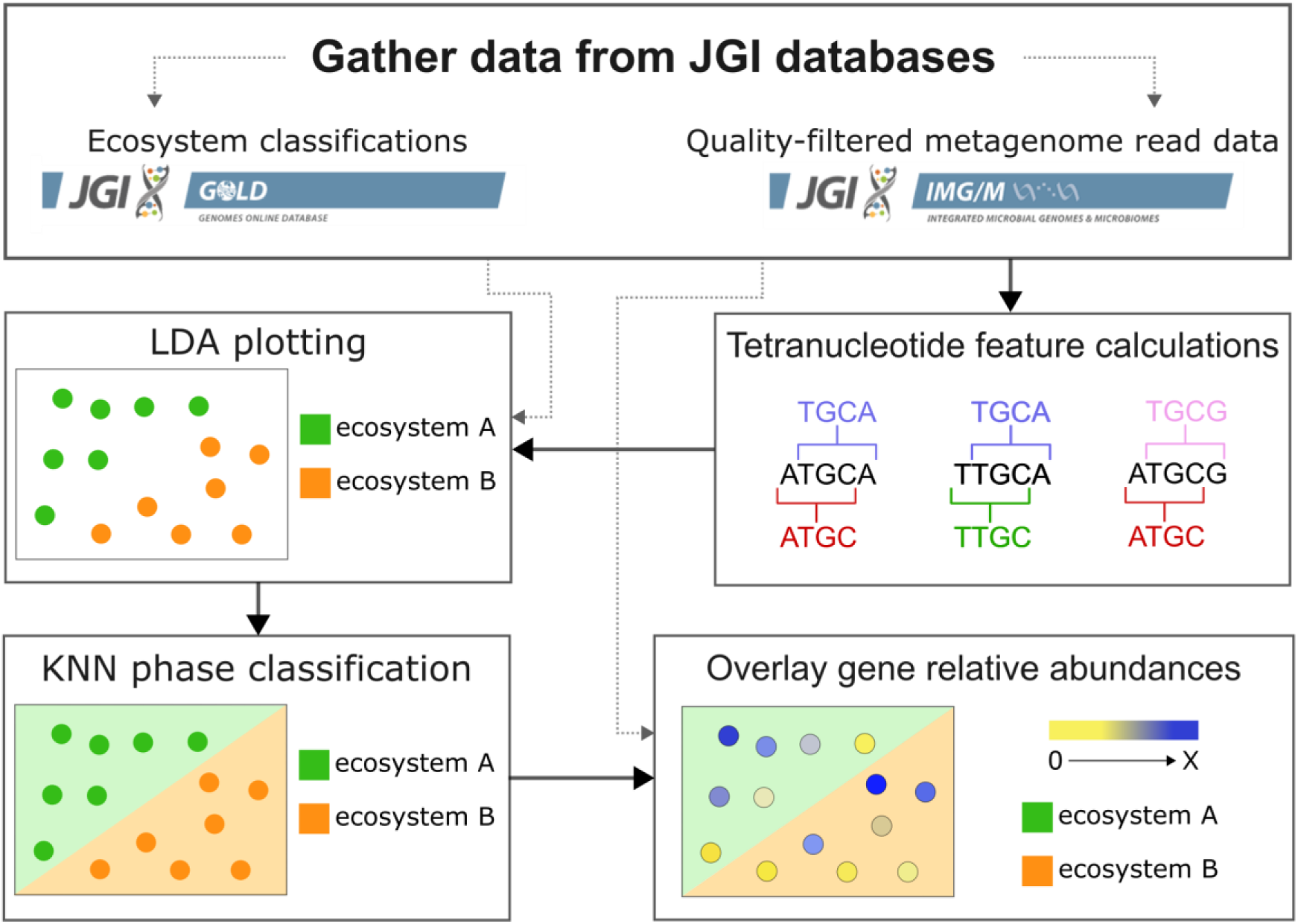
Overview of the analysis workflow to generate the metagenome stability diagram. First, quality-filtered and unassembled sequencing read data from IMG/M metagenomes were combined with ecosystem classifications from GOLD. Tetranucleotide frequencies were calculated for each metagenome, then linear discriminant analysis (LDA) plotting was applied with GOLD ecosystem classification supervision. KNN was applied to LDA coordinates and GOLD ecosystem classifications to map KNN ecosystem phase classifications. Lastly, application of a heatmap overlay for IMG/M gene annotation relative abundances was applied.

Since the number of tetranucleotide features for each metagenome creates far more dimensions than can be visualized (9,316 tetranucleotide frequencies and ratios of frequencies), dimensionality reduction is required for plotting in a two-dimensional space. We used LDA dimensionality reduction to supervise x- and y-coordinate plotting of metagenomes according to both their tetranucleotide composition and GOLD Ecosystem classification labels. LDA seeks to project multidimensional data (tetranucleotide features) into axes that maximize the distance between labeled groups (GOLD Ecosystem classifications) and minimize variation within groups of the same label (Xanthopoulos, Pardalos, and Trafalis 2013). Once metagenomes were plotted with LDA x- and y-coordinates, we used the KNN algorithm to delineate ecosystem classification regions within the two-dimensional space (further described in Methods), which could then be used to evaluate metagenomes sampled along environmental gradients (**Fig. 2**). The resulting interactive metagenome stability diagram is hosted on the IMG/M site as a publicly available resource (https://img.jgi.doe.gov/cgi-bin/mer/main.cgi?section=AdvAnalytics&page=environmental).

**Figure 2.**
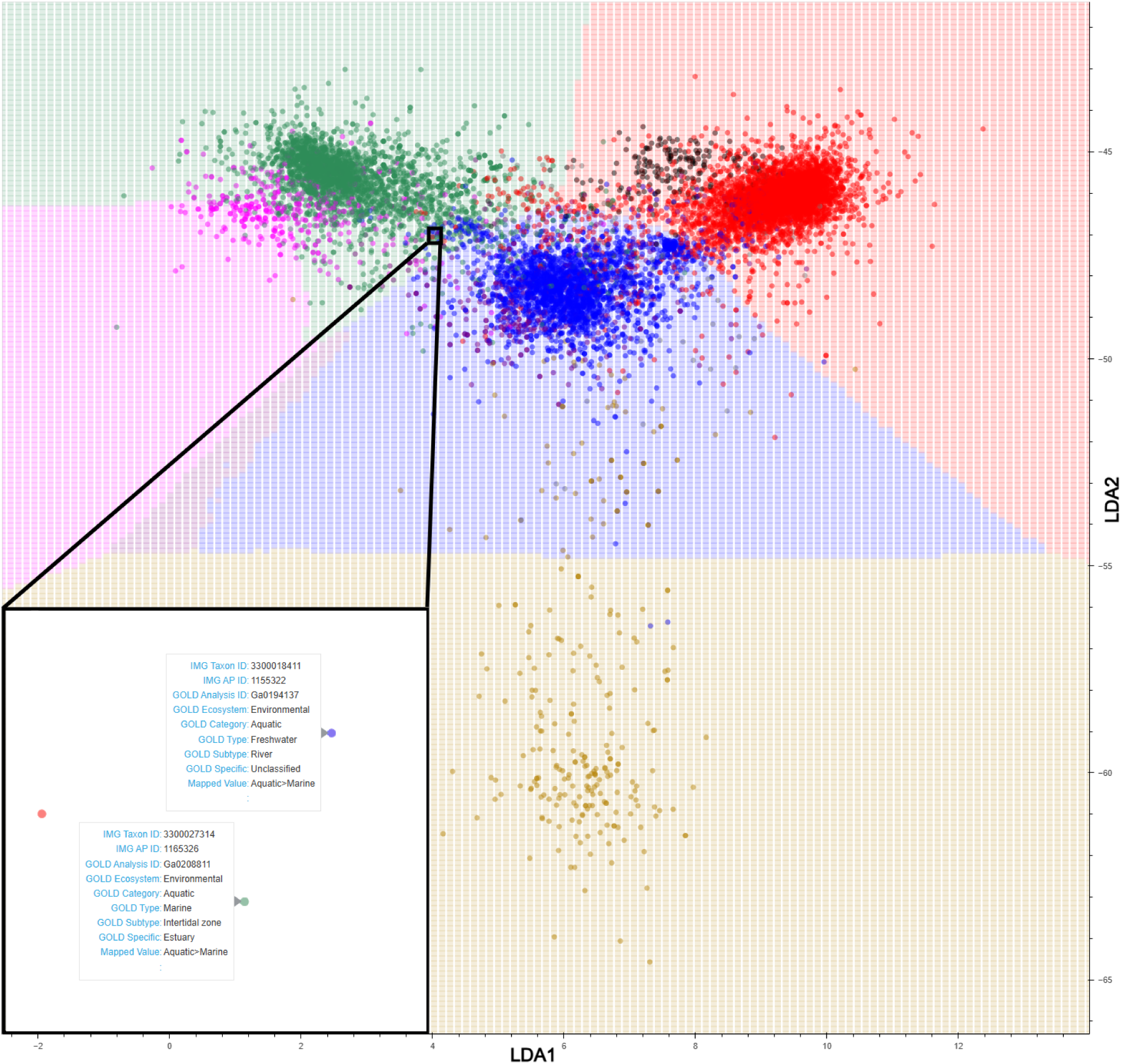
Metagenome stability diagram created with LDA dimensionality reduction of tetranucleotide frequencies and ratios of tetranucleotide frequency and KNN application. Each data point represents a metagenome, colored according to their GOLD Ecosystem Type classification, with the most abundant being Soil (red), Freshwater (blue), Marine (green), Non-marine Saline and Alkaline (pink), Thermal springs (gold), Plant litter (black), Deep subsurface (purple). A full list of metagenomes used is included in **Supplemental File S1**. A zoomed-in panel (bottom left) with IMG and GOLD details is used to demonstrate each point in the diagram as an individual metagenome. Background regions represent KNN Ecosystem classification stability phases of the same color scheme as the metagenome data points, with the exception of Plant litter which does not have a mapped phase due to lack of KNN support. The counts of plotted metagenomes from each KNN Ecosystem classification is available in **Supplemental File S1**. Metagenomes that have a matching GOLD Ecosystem Type data point color and KNN Ecosystem classification phase background color, have tetranucleotide compositions that resemble other metagenomes of the same GOLD Ecosystem Type, as derived by LDA with KNN classification. Metagenomes data points that plot over differing KNN Ecosystem classification phases have tetranucleotide compositions that more closely resemble the differing phase rather than their own GOLD Ecosystem Type classification.

Each data point of the metagenome stability diagram in **Figure 2** represents an individual metagenome and the background shows KNN Ecosystem classification boundaries, with colors representing the metagenomic space of GOLD Ecosystem Type classifications: “Terrestrial>Soil” (red), “Aquatic>Freshwater” (blue), “Aquatic>Marine” (green), “Aquatic>Non-marine Saline and Alkaline” (pink), “Aquatic>Thermal springs” (gold), and “Aquatic>Deep Subsurface” (purple). The KNN Ecosystem classification boundaries between each of these metagenomic phases (e.g. gold to blue meaning “Aquatic>Thermal springs” to “Aquatic>Freshwater”) represent Ecosystem Type transitions, where the tetranucleotide content shifts from a composition resembling metagenomes with one GOLD Ecosystem Type classification to metagenomes of another GOLD Ecosystem Type. The large condensed clusters of data points represent metagenome compositions that are commonly found for the particular KNN Ecosystem classification phase that they are plotted in. These metagenomes remain largely unseparated by LDA projection of within group variation and form a “central” grouping. Metagenome data points that are “peripheral” to the central core metagenomes represent uncommon tetranucleotide compositions, and often originate from lesser sampled GOLD Ecosystem Subtype environments.

### A view into community transitions across ecosystems

Insight into environmental microbiome properties that help to define both central and peripheral metagenome classifications could help reveal microbe-environment interactions that shape our biosphere (Hugerth and Andersson 2017). The transition boundaries between KNN Ecosystem classification phases represent KNN-classification ambiguity, where metagenomes plotted near these boundaries can be labeled with either classification, presumably as environmental conditions approach high similarity. Exploring these transition-associated GOLD Ecosystem classifications more specifically, most of the fringe metagenomes plotted near classification transition boundaries are sampled from environments along ecosystem gradients, such as “Wetlands,” “Estuaries,” and “Sediment,” where two environments mix (e.g. “Wetlands” as a mix between “Soil” and “Freshwater”). The boundary between “Marine” and “Non-marine Saline and Alkaline” classifications in particular appears the least resolved from our collection of metagenomes. Intuitively, these metagenomes would likely be sampled from marine coastal mixing environments, and in fact many are sampled from “Coastal,” “Inlet,” and “Intertidal” GOLD Ecosystem Types where estuarine mixing occurs (Underwood et al. 2022). In **Figure 2**, the KNN Ecosystem classification of an “Aquatic>Deep subsurface” phase did not contain any plotted metagenomes, but presence of a KNN classification phase implies most nearby metagenomes are annotated as such. Indeed, a large number of “Aquatic>Deep subsurface” annotated metagenomes within the “Aquatic>Freshwater” phase are skewed toward the lower left corner, in the direction of the “Aquatic>Deep subsurface” phase. The “Aquatic>Deep subsurface” phase borders the “Aquatic>Marine,” “Aquatic>Non-marine Saline and Alkaline,” “Aquatic>Freshwater,” and “Aquatic>Thermal springs” phases which indicates that any hypothetical metagenomes in the “Aquatic>Deep subsurface” phase would resemble a mixing of microbial communities from those environments.

### Ranked list of protein-coding genes that differentiate across ecosystems

Metagenomes within each KNN Ecosystem classification phase represent different microbial community compositions with differing metabolic capabilities. In addition to GOLD Ecosystem classification annotations, metagenomes have been processed through the IMG annotation pipeline, providing functional information *via* protein-coding gene identifiers. Gene relative abundances, normalized to the amount of assembled bases, are calculated for metagenomes and can be grouped according to KNN Ecosystem classifications of **Figure 2** and/or GOLD Ecosystem classifications. In the IMG/M interactive tool, gene abundances were overlaid on metagenome data points with a heatmap-like color scheme and show patterns of metabolic capabilities in the context of their KNN Ecosystem classification phase. The heatmap color range can be manually adjusted with the “Heat” slider. As an example, **Figure 3a** shows normalized gene abundance heatmap values for COG3237, an uncharacterized conserved protein subunit (YjbJ). When summarized as a box plot (**Fig. 3b**), COG3237 has significantly greater abundances in the “Terrestrial>Soil” KNN Ecosystem classification, suggesting an ecosystem-specific role for this uncharacterized protein.

**Figure 3.**
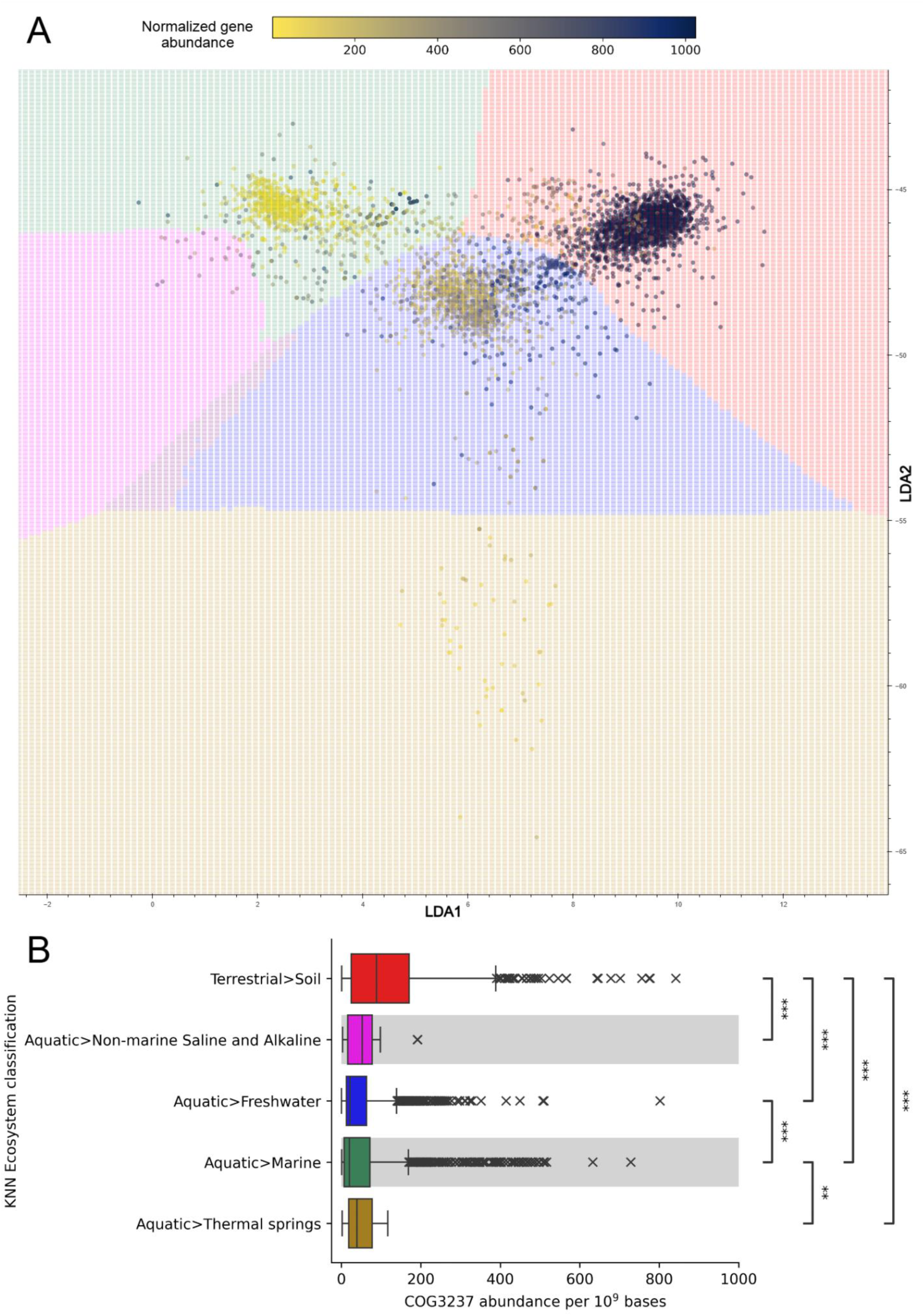
Ecosystem distribution of CO3237 normalized abundance. (a) metagenome stability diagram showing abundances of COG3237, uncharacterized conserved protein YjbJ, normalized per 10^9^ assembled bases. The MatPlotLib (Hunter 2007) reverse cividis colormap was applied to plotted abundances, with a perceptual uniform sequential scale from low abundance (yellow) to high abundance (dark blue). Metagenomes that are missing COG3237 annotation are hidden and not assigned a color. Background color assignment and plotting coordinates are the same as in Figure 1. (b) A boxplot summary of COG3237 normalized abundances shown in 3a. Welch’s ANOVA Games-Howell post-hoc test was used to calculate pairwise statistical significance between non-equal variance KNN Ecosystem classification groups; *p-value≤0.05; **p-value≤0.01; ***p-value≤0.001.

Using a random forest model trained on the metagenome KNN Ecosystem Type classifications and protein-coding annotations, we extracted an ordered list of Clusters of Orthologous Genes (COGs) (Galperin et al. 2021), Enzyme Commission (EC) (McDonald and Tipton 2023), Kyoto Encyclopedia of Genes and Genomes Orthology (KO) (Kanehisa et al. 2024) and European Bioinformatics Institute protein families (Pfams) (Mistry et al. 2021) gene identifiers that exhibit differential abundance across all ecosystems and identifiers that differentiated specific ecosystems from the others (**Supplemental File S3**). The gene identifiers that best differentiate each plotted KNN Ecosystem Type classification, either because they are abundant or sparse, are presented in Table 1. While all of the top differential gene identifiers were COG annotation, gene identifiers EC, KO, and Pfam were included in the analysis as well. These ranked gene identifiers help to confirm the separation of metagenomes based on genomic properties evolved from life-environment interactions, and metabolic properties that confer selective advantage or disadvantage.

**Table 1.** Top gene identifiers that are differentiated by each KNN Ecosystem Type classification and their annotation description. A full list of annotated genes is available in Supplemental File S3.

| KNN Ecosystem Type classification | Most differential gene identifier | Annotation description |
| --- | --- | --- |
| Terrestrial>Soil | COG3237 | Uncharacterized conserved protein YjbJ, UPF0337 family |
| Aquatic>Thermal springs | COG1483 | Predicted ATPase, AAA+ superfamily |
| Aquatic>Non-marine Saline and Alkaline | COG1191 | DNA-directed RNA polymerase specialized sigma subunit FliA |
| Aquatic>Marine | COG4208 | ABC-type sulfate transport system, permease component CysW |
| Aquatic>Freshwater | COG3329 | Na<sup>+</sup>-dependent bicarbonate transporter SbtA |
| Aquatic>Deep subsurface | COG4869 | Propanediol utilization protein PduL |

Three of the six top classification-specific gene identifiers are involved in the acquisition and utilization of nutrients. The prevalence of COG3329 and sparsity of COG4208 were the top genes differentiating “Aquatic>Freshwater” and “Aquatic>Marine” KNN classifications, respectively. The Na^+^-dependent bicarbonate transporter SbtA (COG3329) and the permease component of an ABC-type sulfate transport system CysA (COG4208), are both nutrient transporters that indicate metabolism specificity in response to the environment. The high abundance of *sbtA* in freshwater environments suggests a demand for bicarbonate uptake, which is consistent with the observation of ubiquitous presence of this gene in cyanobacteria genomes often found in freshwater ecosystems (X.-Y. Liu et al. 2021). The cause of low abundance *cysW* in marine environments is more difficult to hypothesize, since low abundances in genomes could be due to alternative transporter presence or a lower need for sulfate uptake transport systems in marine environments (Marietou et al. 2018). Despite containing no metagenome data points plotted within the “Aquatic>Deep subsurface” classification phase, we were able to identify gene identifiers that had abundances most associated with metagenome proximity to the phase. Propanediol utilization protein *pduL* was the top gene for differentiating “Aquatic>Deep subsurface” and is involved in the formation of bacterial metabolosomes for catabolic metabolism of propanediol, a product of sugar fermentation, and may provide a selective advantage in anaerobic environments like the deep subsurface (Yang et al. 2022; Y. Liu et al. 2007).

As previously shown in **Figure 3**, *yjbJ* is highly abundant in soil environments with a function that is largely unknown, but its expression is increased under osmotic stress (Weber, Kögl, and Jung 2006). *yjbJ* was also one of the top gene identifiers to show differential abundance across all ecosystem classifications, with high abundances in soil ecosystems and a gradual decrease across freshwater to low abundances in marine ecosystems (**Fig. 3**), seemingly in agreement with reports of *yjbJ* having a potential role in desiccation resistance (Barcarolo et al. 2020). The predicted ATPase of the AAA+ superfamily, abundant in thermal spring environments; and fliA, abundant in non-marine saline environments, are less specific toward metabolic processes and instead may be indicative of highly abundant taxa and/or taxa that contain many copies of these genes. While all six top classification-specific gene identifiers were COG identifiers, the ranked lists include all gene identifiers used in the random forest model.

#### Investigating metabolic interactions with the environment

The use of protein annotations in a metagenome stability diagram enables visualization of successful metabolic strategies. Using protein-coding annotations to estimate environmental nutrient concentrations and speculate on nutrient/energy capture is established in microbial ecology (Kellom et al. 2022; Reji et al. 2019; Hamilton et al. 2019; Gutiérrez-Preciado et al. 2018). For certain substrates, cells must maintain the proper balance of flux to meet metabolic demands while also avoiding toxicity (Müller et al. 2006). Copper is a prime example, because as a metal cofactor it is required for many enzymes to function while also being toxic if internal concentrations become too great (C. Li, Li, and Ding 2019). The distribution of *ycnJ* (KO:K14166), a copper importer protein subunit (Chillappagari et al. 2009) and *copC* (KO:K07156), a copper-resistance protein subunit (Raimunda et al. 2011; Lawton et al. 2016; Chillappagari et al. 2009) across “Aquatic>Freshwater” and “Terrestrial>Soil” KNN Ecosystem classification phases highlights differing metabolic strategies for copper flux balance within each environment (**Fig. 4**). These two genes encode for proteins contained within the same COG identifier (COG2372) and highlight the complicated and understudied mechanisms of copper homeostasis. YcnJ is a homolog of the CopCD dimer and functions as a copper importer in genus *Bacillus* (Chillappagari et al. 2009; B. Li et al. 2024), and CopC has an unclear mechanism for imparting copper resistance with potential roles in copper efflux, sequestration, and import (Lawton et al. 2016; Udagedara et al. 2019). While these two genes do not provide a complete view of copper homeostasis (Solioz 2018), **Figure 4** shows the prevalence of potential copper resistance *via* CopC relative to YcnJ-mediated copper influx for specific GOLD Ecosystem Subtypes split across two KNN Ecosystem classification phases. In the “Aquatic>Freshwater” phase, higher abundances of *copC* are found in Lake, River, and Groundwater subtypes, suggesting microbial populations that are susceptible to copper-toxicity and possibly exist in copper-replete conditions, where copper-resistance is beneficial. GOLD Ecosystem Subtypes of Warm (34-46C), Tepid (25-34C), and Intertidal zone, had higher abundances of *ycnJ*, indicating a demand for copper among those populations, which likely includes the genus *Bacillus*. Despite relatively high counts of metagenomes within the “Aquatic>Freshwater” phase, Wetlands, Sediment, and Unclassified GOLD Ecosystem Subtypes had no difference between *copC* and *ycnJ* abundances. In the “Terrestrial>Soil” phase, all GOLD Ecosystem Subtypes with a greater number of metagenome samples than in the “Aquatic>Freshwater” phase had higher abundances of *ycnJ*, except Peat, Glacial till, and Temperate forest. Higher abundances of *ycnJ* suggest that soil populations are generally in need of copper supply. Conversely, Glacial till and Peat had higher abundances of *copC*, and along with the Lake, River, and Groundwater subtypes of the “Aquatic>Freshwater” phase, may contain populations that benefit from copper-resistance. Removing or adding copper in these ecosystems would likely create instability for these microbiomes, triggering a metagenome compositional shift (Thomas et al. 2023) toward a nearby region of their classification phase that has tolerable copper conditions.

**Figure 4.**
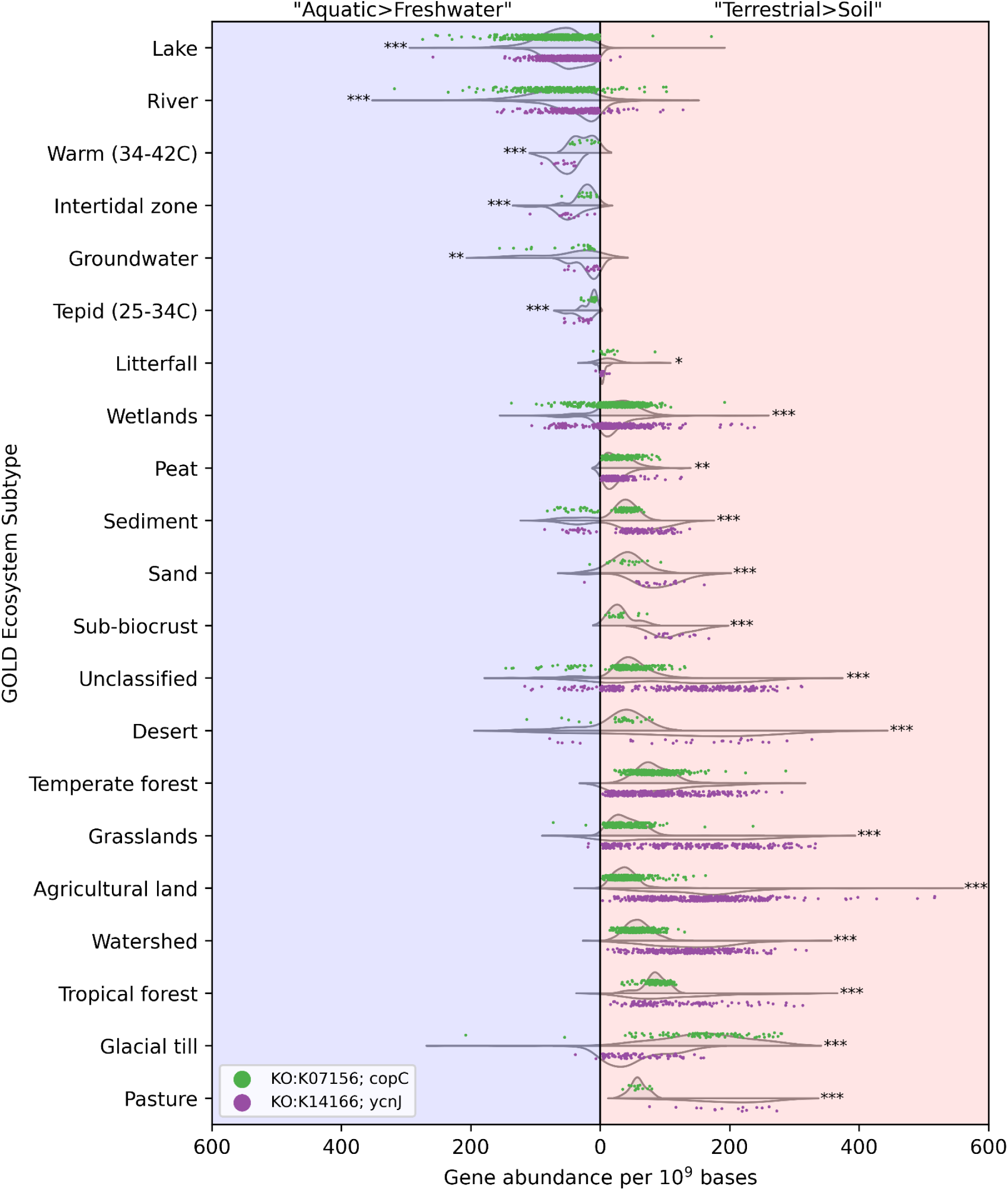
The distribution of KO:K07156 (copC) (green points) and KO:K14166, (ycnJ) (purple points) gene abundances relative to 10^9^ assembled bases across GOLD Ecosystem Subtypes within the “Aquatic>Freshwater” (blue background, left of vertical line at x=0) and “Terrestrial>Soil” (red background, right of vertical line at x=0) metagenome stability diagram KNN Ecosystem classification phases of Figure 2. Metagenomes with no abundance data available for either gene are not included in the plot. Only GOLD Ecosystem Subtypes with >10 representative metagenomes are included, totalling 3,865 metagenomes across 21 Subtypes. Split violin plots highlight data point density for KO:K07156, (*copC*) (top-side violin) and KO:K14166, (*ycnJ*) (bottom-side violin) within each GOLD Ecosystem Subtype. Two-tailed t-test statistical significance of each KO:K07156, (*copC*) and KO:K14166; (*ycnJ*) pairing was calculated for “Aquatic>Freshwater” and “Terrestrial>Soil” data points separately; *p-value≤0.05; **p-value≤0.01; ***p-value≤0.001.

We also used protein annotations of metagenomes to decipher patterns of nutrient and elemental cycling across ecosystem classifications. The nitrogen cycle, which includes both oxidative and reductive pathways along with assimilatory and dissimilatory nitrogen usage (Kuypers, Marchant, and Kartal 2018), is extensively studied based on metagenomic data (Zhang, Ward, and Sigman 2020; Hutchins and Capone 2022; Santoro et al. 2017). Nitrate reduction to nitrite is the first step of multiple reducing pathways with differing end-products, each with their own microbiome implications. **Figure 5** explores the metagenomic patterns of nitrite reduction, looking at the fate of nitrite in the global nitrogen cycle. *nirK* (KO:K00368) is a gene marker for copper-containing nitric oxide forming nitrite reductase, involved in a process called denitrification in anaerobic environments, reducing nitrite to gaseous nitrous oxide (Graf, Jones, and Hallin 2014). Denitrification will eventually produce N_2_O and N_2_ gas, which is no longer a fixed-nitrogen source and lost to the atmosphere. *nirK* abundances were highest in soil metagenomes with sparse abundance peaks in aquatic environments having high sediment contact, such as rivers, floodplains, intertidal zones, and hydrothermal vents, which may represent anaerobic conditions (**Supplemental File S4**). *nrfA* (KO:K03385) is a gene marker for heme-containing dissimilatory nitrite reductase that reduces nitrite to ammonium, using nitrite as an electron acceptor and keeping nitrogen fixed in the environment (Besson, Almeida, and Silveira 2022). In addition to nitrogen-fixation and organic nitrogen catabolism, dissimilatory nitrite reduction supplies bioavailable ammonium to the environment, which feeds nitrogen metabolism in oxidative conditions. *nrfA* abundances were generally highest in thermal spring environments with abundance peaks in restoration grassland soil environments with agricultural history, although the highest abundance is from a Antarctica dry valley desert soil metagenome. In the case of restoration grassland soil metagenomes, the high abundance of *nrfA* is likely due to the regrowth of native perennial leguminous plants after fertilizer soil amendments have ceased (Mason et al. 2023), which selects for symbiotic nitrogen-fixing bacteria. Assimilatory nitrate reductase marker *nirB* (KO:K00362) also converts nitrite to ammonium, which is then in turn used for the synthesis of biomolecules (Besson, Almeida, and Silveira 2022). *nirB* abundances were also highest in soil environments, but more prevalent across aquatic environments than *nirK* and *nrfA*. Grouping metagenomes by genomic content and highlighting metabolic capabilities helps to decipher the fate of global nutrient cycling across ecosystem classifications. In the case of nitrite, in **Figure 5**, we have observed metabolic abundance patterns that drive the bioavailability of nitrogen as it cycles globally.

**Figure 5.**
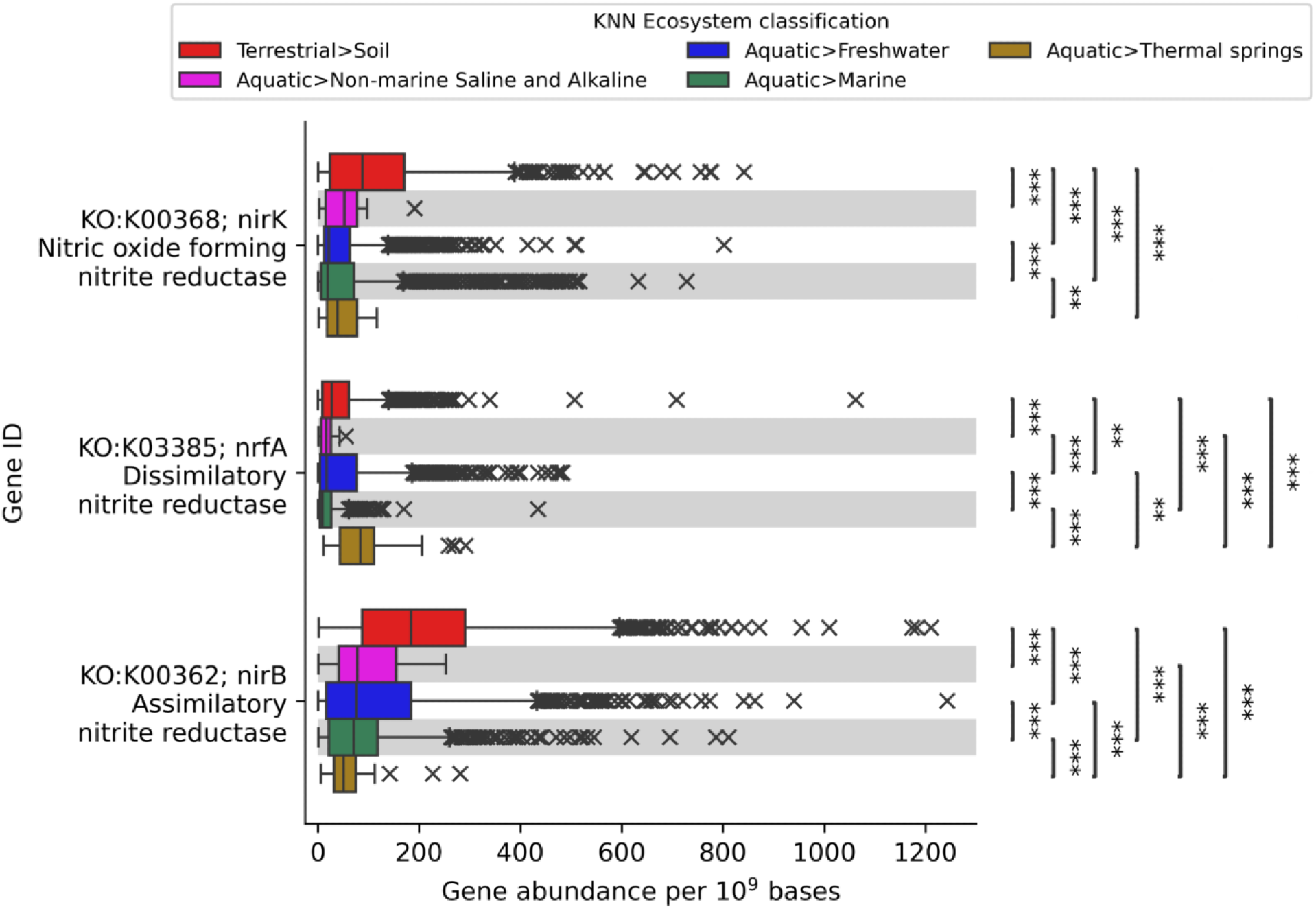
Nitrite reductase box plot showing normalized gene abundances for KO:K00368 (nirK), KO:K03385 (nrfA), and KO:K00362 (nirB) across KNN Ecosystem classification metagenomes of the metagenome stability diagram of Figure 2. Welch’s ANOVA Games-Howell post-hoc test was used to calculate pairwise statistical significance between non-equal variance KNN Ecosystem classification groups; *p-value≤0.05; **p-value≤0.01; ***p-value≤0.001.

## Discussion

This work aims to determine genome-derived boundaries that separate metagenome ecosystem classifications and help reveal microbe-environment interactions that shape our biosphere. We use tetranucleotide compositions and ecosystem classification observations to delineate boundaries between metagenome groups, similar to the use of chemical compositions delineating mineral samples in mineral stability diagrams (Faure 1998). Tetranucleotides are universally shared components of (meta)genomes (Noble, Citek, and Ogunseitan 1998), which means that any distance between metagenomes within our metagenome stability diagram is a result of varying tetranucleotide frequencies. Metagenome samples can be thought of as aggregates of microbial populations, and since tetranucleotide frequencies are taxonomy specific (Pride et al. 2003; Teeling et al. 2004), microbial population composition is the determining factor for metagenome coordinates within **Figure 2**. In contrast, other k-mer-based implementations of determining metagenomic distance can be supported by comparing their fraction of shared k-mers (Ondov et al. 2016; Pierce et al. 2019). Although both methods of determining metagenomic distance are based on microbial community composition, the distance between hypothetical metagenomes with no shared k-mers is incalculable, which is increasingly possible with increasing k-mer size and taxonomic distance (Bussi, Kapon, and Reich 2021). While the small size of tetranucleotides often decreases taxonomic specificity for identification when compared to larger k-mers (Bussi, Kapon, and Reich 2021), microbial community taxonomy is not needed for our purposes of metagenome plotting.

One assumption of our analysis is that each metagenome represents a stable microbiome in its plotted KNN Ecosystem classification phase, i.e. having a microbial composition that is adapted to its environmental conditions. Within each classification phase however, multiple variations of microbiome stability can exist, since each microbial community has a past history of adaptation to niche perturbations (Shade et al. 2013; Stegen, Bottos, and Jansson 2018; Correa-Garcia, Constant, and Yergeau 2022; Gonze et al. 2017; Stein et al. 2013; Nuccio et al. 2020). This contributes to the clustering spread that can be observed in all classification phases. Interestingly, the “Aquatic>Thermal springs” classification phase is relatively less clustered than all others, which could be due to the heterogeneous environments classified as thermal springs and the range of potential metabolisms (Shock et al. 2010; Alsop, Boyd, and Raymond 2014). Metagenoms are also less clustered near KNN Ecosystem classification phase transitions, which may be representative of varying environmental conditions along a gradient between GOLD Ecosystem Types. While phase transitions represent environment mixing, the physicochemical properties are constant enough to drive the formation of stable microbial compositions (J. M. Dick et al. 2023; Gonze et al. 2017). Although the “Aquatic>Deep subsurface” phase currently lacks metagenome data points, as more metagenomes are plotted into the metagenome stability diagram, the coordinates of this phase may migrate to a region honing in on tetranucleotide composition distinction from “Aquatic>Freshwater” that reflects environmental physicochemical properties. Similarly, “Terrestrial>Plant litter” annotated metagenomes appear to be nearing tetranucleotide composition distinction from “Terrestrial>Soil” metagenomes, but have not yet reached a density within the **Figure 2** LDA coordinates such that KNN classification of a “Terrestrial>Plant litter” phase has been plotted. Areas of the metagenome stability diagram that do not yet contain plotted metagenomes represent hypothetical microbial community compositions from niche environments that have yet to be sampled. Taking into account metagenomes that are nearby on the metagenome stability diagram, areas that are so far unrepresented by data points may guide future sampling expeditions.

Recently, the clustering of KO gene identifiers from IMG/M metagenomes identified metagenome clusters reflecting terrestrial, aquatic, and anaerobic ecosystems (Flinkstrom et al. 2024). As part of their analysis, three clusters were differentiated by functional markers and GC content. Remarkably, their study which clustered metagenomes by protein-coding annotations and resulted in a sequence content differentiation, while the opposite of our approach clustering with unassembled sequence content (tetranucleotides from quality-filtered sequence reads), identified similar top ecosystem-specific differentiating gene markers (**Supplemental File S3**). Across all ecosystems, Flinkstrom and colleagues found two non-homologous end-joining DNA repair genes positively correlated with their finding of GC content differentiation among clustered metagenomes, *ligD* and *ku*. From our analysis, the *ku* gene identifier COG1273 was the fourth highest overall differentiating gene identifier, with KO:K10979 (KEGG gene identifier for *ku* as in (Flinkstrom et al. 2024)) as the 514^th^ highest, out of 45,536 total gene identifiers. *ligD*, as KO:K01971, was the 336^th^ highest gene identifier overall in our analysis, and 283^rd^ as COG1793. The ranking of gene identifiers as differentiating across and within ecosystem classification could be an important guiding tool for future studies characterizing proteins of unknown function or refining concepts of protein function and metabolism dispersal.

## Conclusion

Metagenome-ecosystem stability implies that microbial populations are able to acquire, metabolize, and excrete chemical compounds required for growth, to which their lineage has become adapted (Jansson and Hofmockel 2020). Understanding these processes relies on comparison of both biological and environmental components (Swingley et al. 2012; Alsop, Boyd, and Raymond 2014). Any environmental conditions that limit metabolic processes help to create ecosystem classification boundaries within our metagenome stability diagram, since microbial composition will be forced to adapt (Okie et al. 2020; Moore et al. 2013). There is also an expectation that if nutrients exist with favorable environmental conditions, a microbial population will grow and consume those nutrients (Zwetsloot et al. 2020; Lee et al. 2017). With this in mind, the metabolic potential of metagenomes in the form of protein-coding annotations provides insight on environmental conditions and nutrient flux. Future metagenomes sequenced, assembled, and annotated by the Joint Genome Institute will be analyzed with the methods laid out here and contribute to future annual updates to the interactive metagenome stability diagram tool in IMG/M. This work could help lead to a future of microbiome science where microbial population compositions can be predicted based on initial and/or perturbed environmental conditions, and possibly have implications for microbiome engineering.

## Methods

Publicly available metagenomes sequenced at the JGI and added to the IMG database before April 10^th^ 2024 (date of collection) were considered for this analysis (Chen et al. 2023). This data collection criteria yielded 15,208 metagenome datasets labeled as “Metagenome Analysis” as their GOLD Analysis Project Type (Mukherjee et al. 2024). Metagenomes that had a GOLD Ecosystem classification of “Engineered” or “Host-associated” or had a GOLD Ecosystem Type classification of “Nest” were removed from our metagenome collection for non-natural or host-restricted environment properties that would alter the interpretation of tetranucleotide frequencies plots as reflecting physicochemical pressures. After data filtering, the total number of included metagenomes was 12,063, with GOLD Ecosystem Categories and Types can be found in **Supplemental File S1**.

From the included metagenome datasets, filtered sequencing reads following the standard operating procedure of the DOE-JGI Metagenome Annotation Pipeline (Huntemann et al. 2016) were analyzed for tetranucleotide counts with KMC (version 3.1.1) (Kokot, Długosz, and Deorowicz 2017). Tetranucleotide counts were converted into frequencies (e.g. 4-mer count divided by the total count of all 4-mers) and ratios of frequencies (e.g. 4-mer frequencies divided by all other permutations of 4-mer frequencies) (**Supplemental File S2**). Tetranucleotide frequencies and ratios of frequencies were used as features for linear discriminant analysis (LDA) dimensionality reduction using Scikit-learn (v1.5.2) (Pedregosa et al. 2011) to plot metagenomes in a two-dimensional space with reference to their GOLD Ecosystem Type classification. Including the ratios of tetranucleotide frequencies as features for our metagenome LDA dimensionality reduction allows additional vector coefficients (Zhao et al. 2020) to be utilized for roughly estimating relative abundances between taxa/metabolisms. LDA is a supervised algorithm that calculates x- and y-coordinates that maximize group separation and minimizes intra-class variation (Zhao et al. 2020). Metagenome groups were defined as “GOLD Ecosystem Category>GOLD Ecosystem Type” (**Supplemental File S1**). Once metagenome x- and y- coordinates were plotted, k-nearest neighbors (KNN) was used to define metagenome stability phases as KNN Ecosystems classifications. A value of k=500 nearest metagenomes was chosen based on diminishing accuracy, precision, and F1-score statistics when increasing k, using a training/testing data split of 80%/20% (**Supplemental Figure S1**). Values of k<500 showed evidence of overfitting with GOLD Ecosystem Type classification boundaries deviating to include outlier data points. Overfitting at lower k values is not surprising since KNN as a classifier is prone to overfitting due to noise-sensitivity at lower values of k, and can be ameliorated by employing larger k values (Ougiaroglou and Evangelidis 2015). The metagenome stability phases that are plotted with KNN in **Figure 2** are: “Terrestrial>Soil”, “Aquatic>Freshwater”, “Aquatic>Marine,” “Aquatic>Non-marine Saline and Alkaline,” “Aquatic>Thermal springs,” and “Aquatic>Deep Subsurface.” Data visualization was inspired by (Kaushik 2019) showing how to visualize classifier decision boundaries. Visualization was done with the Python-based customizable plotting tool Bokeh (v3.4.3) (Bokeh Development Team 2024).

Protein-coding annotation identifiers (COG, EC, KO, PFAM) were counted for metagenomes with gff3-format annotation files resulting from the IMG annotation pipeline v.5.0.0 or newer, which was 5,683 out of the total 12,063 plotted metagenomes. These annotation counts were normalized to the total number of assembled bases for each metagenome, and used as heatmap “Map Values” to be overlaid on top of metagenome data points. Statistical tests were calculated with the python package Pingouin (Vallat 2018). In the interactive tool, each heatmap “Map Value” was multiplied by a constant of 10^9^ to yield an appropriate MatPlotLib (Hunter 2007) reverse cividis colormap value implemented in Bokeh. Metagenomes without gff3-format annotation files were assigned a heatmap “Map Value” of “NaN” and are hidden from the protein annotation view of the interactive tool. Since protein-coding annotations can have a wide range of counts depending on the gene, the interactive tool “Heat” slider can adjust the heatmap color values median to better illuminate ubiquitously lower or higher counts.

With Scikit-learn’s RandomForestClassifier, we used a random forest model with a 75%/25% training/testing data split on the metagenome KNN Ecosystem Type classifications and protein-coding annotations to extract an ordered list of gene annotation importances according to mean decrease in impurity (Han, Guo, and Yu 2016) (**Supplemental File S3**). These ordered lists were used to identify top KNN Ecosystem classification-specific gene annotations.

## Acknowledgements

The work conducted by the U.S. Department of Energy Joint Genome Institute (https://ror.org/04xm1d337), a DOE Office of Science User Facility, is supported by the Office of Science of the U.S. Department of Energy operated under Contract No. DE-AC02-05CH11231. This research used resources of the National Energy Research Scientific Computing Center (NERSC) (https://ror.org/05v3mvq14), a Department of Energy Office of Science User Facility under Contract No. DE-AC02-05CH11231.

## Author contributions

The study was conceived and designed by M.K. and E.A.E-F. Metagenome data processing and curation was performed by M.K., I-M.A.C., A.C., M.H., N.N.I., S.M., T.B.K.R., S.R., R.S., N.J.V. Formal analysis was performed by M.K. Visualizations were created by M.K. and M.B. Web development was performed by M.K. and K.C. The original draft of the manuscript was written by M.K. and E.A.E-F, with contributions from N.C.K., G.S., and T.W. All authors contributed to the reviewing and editing of the final version of the manuscript.

## Supplemental Material

**Supplemental File S1 - Supplemental_File_S1.xlsx**

Metagenome data table downloaded from the metagenome stability diagram tool, including IMG Taxon IDs, LDA coordinates, plotting colors, KNN Ecosystem classifications (map_classification), IMG Analysis Project IDs, GOLD Analysis IDs, GOLD Ecosystem classifications. A metagenome count summary for each GOLD Ecosystem and KNN Ecosystem classification is included.

**Supplemental File S2 - Supplemental_File_S2.csv**

Tetranucleotide frequencies and ratios of tetranucleotide frequencies of every metagenome included in the metagenome stability diagram tool. Available at https://doi.org/10.5061/dryad.tb2rbp0c8.

**Supplemental File S3 - Supplemental_File_S3.xlsx**

Ranked gene identifier random forest importances as calculated by mean decrease in impurity for all KNN Ecosystem classification as well as overall across all metagenomes.

**Supplemental File S4 - Supplemental_File_S4.xlsx**

Metagenome data table downloaded from the metagenome stability diagram tool, including IMG Taxon IDs, Normalized gene abundance values (map_value), LDA coordinates, plotting colors, KNN Ecosystem classifications (map_classification), IMG Analysis Project IDs, GOLD Analysis IDs, GOLD Ecosystem classifications for “KO:K00368; *nirK*”, “KO:K03385; *nrfA*”, and “KO:K00362; *nirB*” gene abundances in **Figure 5**.

**Supplemental Figure S1 - Supplemental_Figure_S1.pdf**

Supplemental figure depicting the effect of KNN k value on Scikit-learn classification report metrics precision, recall, and F1 scores, with and without the inclusion of ratios of tetranucleotides in the data set.

